# Polled Digital Cell Sorter (p-DCS): Automatic identification of hematological cell types from single cell RNA-sequencing clusters

**DOI:** 10.1101/539833

**Authors:** Sergii Domanskyi, Anthony Szedlak, Nathaniel T Hawkins, Jiayin Wang, Giovanni Paternostro, Carlo Piermarocchi

**Affiliations:** Department of Physics and Astronomy, Michigan State University, 48824 East Lansing, MI, USA.; Salgomed, Inc., 92014 Del Mar, CA, USA.; Sanford Burnham Prebys Medical Research Institute, 92037 La Jolla, CA, USA.

**Keywords:** Single cell RNA sequencing, Cell type identification, Biomarkers, Bone Marrow

## Abstract

**Background:** Single cell RNA sequencing (scRNA-seq) brings unprecedented opportunities for mapping the heterogeneity of complex cellular environments such as bone marrow, and provides insight into many cellular processes. Single cell RNA-seq, however, has a far larger fraction of missing data reported as zeros (dropouts) than traditional bulk RNA-seq. This makes difficult not only the clustering of cells, but also the assignment of the resulting clusters into predefined cell types based on known molecular signatures, such as the expression of characteristic cell surface markers.

**Results:** We present a computational tool for processing single cell RNA-seq data that uses a voting algorithm to identify cells based on approval votes received by known molecular markers. Using a stochastic procedure that accounts for biases due to dropout errors and imbalances in the number of known molecular signatures for different cell types, the method computes the statistical significance of the final approval score and automatically assigns a cell type to clusters without an expert curator. We demonstrate the utility of the tool in the analysis of eight samples of bone marrow from the Human Cell Atlas. The tool provides a systematic identification of cell types in bone marrow based on a recently-published manually-curated cell marker database [1], and incorporates a suite of visualization tools that can be overlaid on a t-SNE representation. The software is freely available as a python package at https://github.com/sdomanskyi/DigitalCellSorter

**Conclusions:** This methodology assures that extensive marker to cell type matching information is taken into account in a systematic way when assigning cell clusters to cell types. Moreover, the method allows for a high throughput processing of multiple scRNA-seq datasets, since it does not involve an expert curator, and it can be applied recursively to obtain cell sub-types. The software is designed to allow the user to substitute the marker to cell type matching information and apply the methodology to different cellular environments.

## Background

Bulk RNA-sequencing has provided the bioinformatics community with a large volume of high quality data over the past decade. However, bulk measurements make studying the transcriptomics of heterogeneous cell populations difficult and provides limited insight on complex systems composed of interacting cell types. Single cell RNA-seq (scRNA-seq) techniques promise to provide the field of bioinformatics with samples suf-ficiently large to resolve the subtleties of heterogeneous cell populations. [2, 3]

The identification of cell types based on specific molecular signatures is challenging. This is particularly true in samples obtained from *ex vivo* bone marrow or periferal blood samples, where different types of hematological cells coexist and interact. scRNA-seq of periferal blood mono-nuclear cells (PBMC) and bone marrow mono-nuclear cells (BMMC) is nowadays possible with high level of sensitivity (see e.g. [4]). Monitoring different cell types and their heterogeneity in these hematological tissues has important applications in precision immunology, and it could help in deter-mining the optimal therapeutic solutions in different hematological cancers.

The classification of the hematopoietic and immune system is predominantly based on a group of cell surface molecular markers named *Clusters of Differentiation* (CD), which are widely used in clinical research for diagnosis and for monitoring disease [5]. These CD markers can play a central role in the mediation of signals between the cells and their environment. The presence of different CD markers may therefore be associated with different biological functions and with different cell types. More recently, these CD markers have been integrated in comprehensive databases that also include intra-cellular markers. An example is provided by CellMarker [1], which will be used here. This comprehensive database was created by a curated search through PubMed and numerous companies’ marker handbooks including R&D Systems, BioLegend (Cell Markers), BD Biosciences (CD Marker Handbook), Abcam (Guide to Human CD antigens), Invitrogen ThermoFisher Scientific (Immune Cell Guide), and eBioscience ThermoFisher Scientific (Cytokine Atlas). However, using these markers on each single cell RNA-seq data for a one-by-one identification would not work for most of the cells. This is fundamentally due to two reasons: (1) The presence of a marker on the cell surface is only loosely associated to the mRNA expression of the associated gene, and (2) single cell RNA-sequencing is particularly prone to dropout errors (i.e. genes are not detected even if they are actually expressed).

The first step to address these limitations is unsupervised clustering. After clustering, one can look at the average expression of markers to identify the clusters. Several clustering methods have been recently used for clustering single cell data (for recent reviews see [6, 7]). Some new methods are able to distinguish between dropout zeros from true zeros (due to the fact that a marker or its mRNA is not present) [8], which has been shown to improve the biological significance of the clustering. However, once the clusters are obtained, the cell type identification is typically assigned manually by an expert using a few known markers [9, 4]. While in some cases a single marker is sufficient to identify a cell type, in most cases human experts have to consider the expression of multiple markers and the final call is based on their personal empirical judgment.

An example where a correct cell type assignment requires the analysis of multiple markers is shown in Fig. 1, where we analyzed single cell data from the bone marrow of the first donor from the HCA (Human Cell Atlas) preview dataset [10] using t-distributed Stochastic Neighbor Embedding (t-SNE) layouts. After clustering (Fig. 1 (a)), the pattern of CD4 expression (Fig. 1 (b)) suggests that cluster 1 (red) and cluster 2 (light green) are both highly enriched for CD4+, potentially indicating T helper cells. In these cells, the expression of CD4 is crucial for sending signals to other types of cells and they are often just called CD4 cells. However, a more careful analysis of cluster 2 shows a significant expression of CD14, CD33 and CD52 (Fig. 1 (c-e)) that indicates that this cluster consists more likely of Macrophages/Monocyte cells.

**Figure 1.**
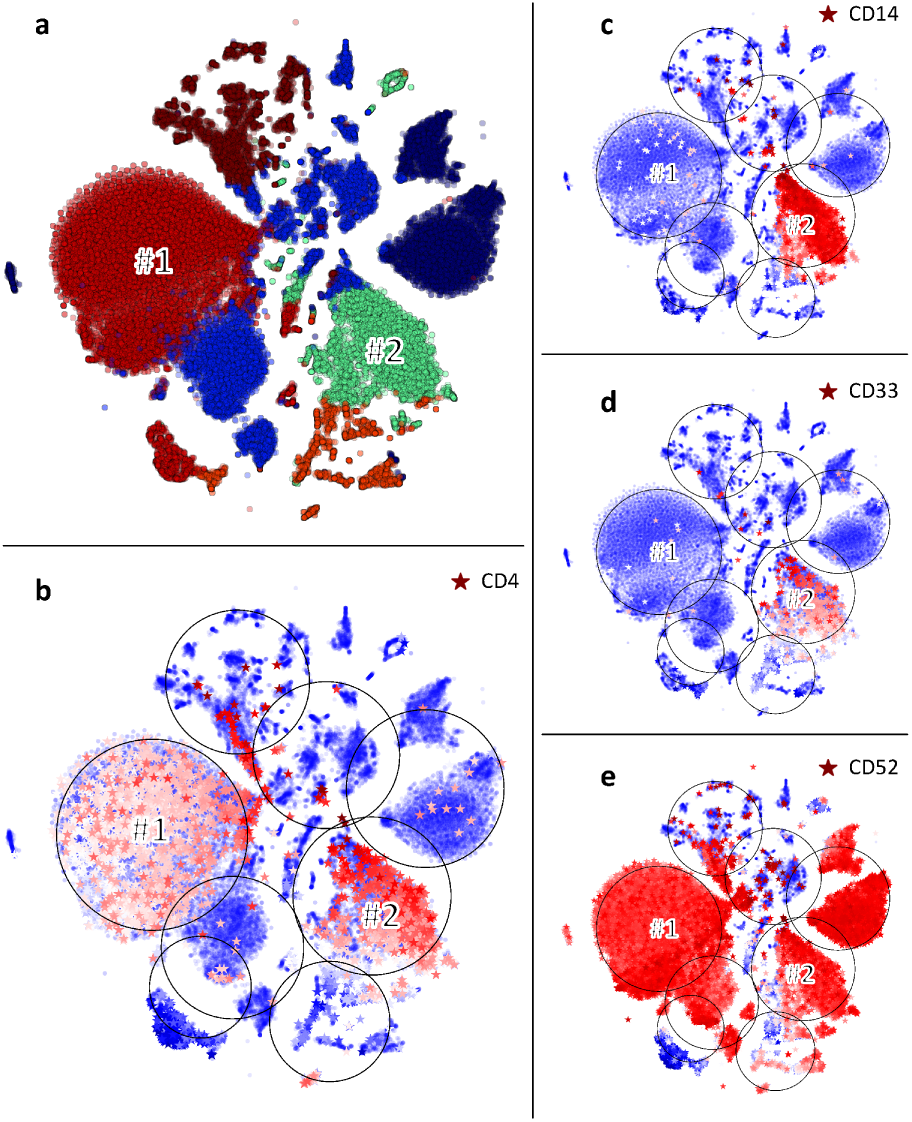
Markers analysis. (a) t-SNE layout of clusters obtained from the first donor of the HCA preview dataset [10]. CD4 marker expression displayed on a t-SNE layout: cells where CD4 is expressed are shown as stars colored according to the expression level from blue (lowest expression) to red (highest expression), large black circles infer the cluster sizes. Cells in which the marker is not expressed are shown as circles. (c-e) Expression of CD14, a myeloid marker, CD33 and CD52 shown as in (b).

In this paper we present a methodology that, after unsupervised clustering, automatically assigns clusters to cell type based on a systematic, unbiased, voting algorithm. Our method does not rely on a human expert empirically selecting a set of markers to interpret the results, but uses all the information available in a large markers database to predict cell types. While cell type identification by manual interpretation can provide good results, the proposed methodology assures that all the available information is taken into account in an unbiased way, and it allows for the identification of many datasets in parallel. From an algorithmic point of view, voting algorithms are among the simplest and most successful approaches to implement fault tolerance and obtain reliable data from multiple unreliable channels [11]. The idea can be traced back to von Neumann [12], and since then it has been practically used in many error correction computational architectures. The voting algorithm employed here belongs to the class of approval voting algorithms. For a given cluster, each participant (a cell marker) votes for a subset of candidates (cell types) that meet the participant criteria (significant RNA expression) for the position rather than picking just one candidate. The approval vote tally determines the score that we use to assign the cluster to a cell type.

## Methods

### Overview

Our p-DCS consists of two main modules: (a) clustering and (b) cell type assignment, which are both based on an unsupervised approach. We demonstrate our methodology using public bone marrow scRNA-seq data from eight donors [10], that will be referred to as BM1-BM8. In this section, we will illustrate the methodology using the first dataset BM1. The remaining bone marrow data along with a large scRNA-seq PBMC dataset, obtained from a different study [4], are analyzed in sec. Results and discussion. In sec. Results and discussion we also show how the proposed methodology can be used recursively, so that for each major cell type one can find the corresponding subtypes. Fig. 2 shows the workflow of the methodology.

**Figure 2.**
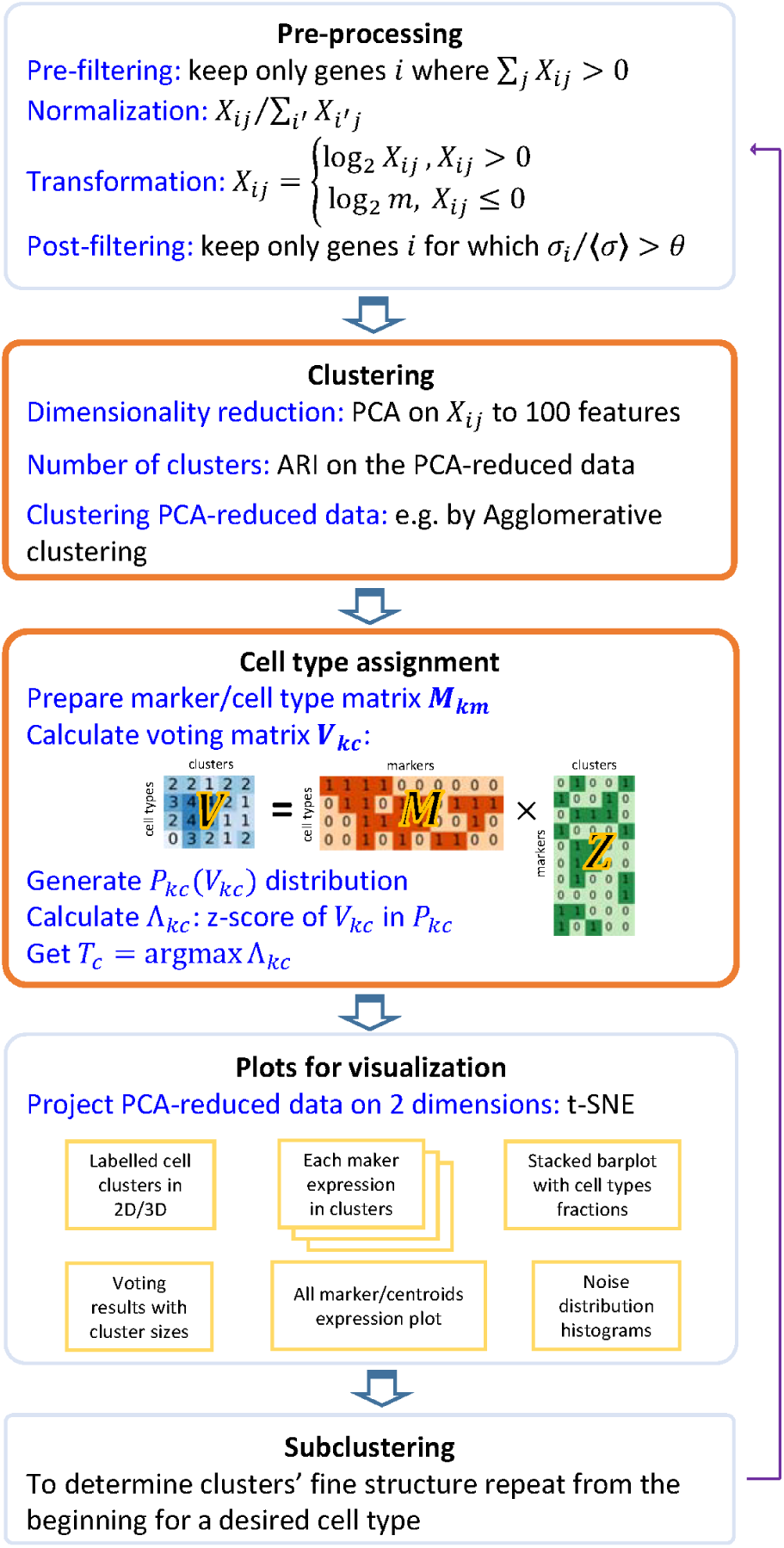
Algorithm schematic. Illustration of the methodology with the two main modules highlighted. The novel polling algorithm for cell identification is implemented in the second highlighted module.

The two main modules are identified by the “Clustering” and “Cell type assignment” labels. The clustering module is preceded by data pre-processing, and a set of visualization tools is included in the software.

### Initial gene/cell filtering and normalization

The expression matrix, *X*_*ij*_, the expression of gene *i* in cell *j* where *i* = 1*, …, N* and *j* = 1*, …, p* is normalized following the steps outlined in [4]. The gene expression matrix is first filtered to keep only genes *i* that are expressed in at least one cell (Σ_*j*_*X*_*ij*_*>* 0). The expression in all cells must then be mapped to the same range of total expression to account for differing yields from PCR amplification. Each cell’s expression vector is thus divided by the sum of all its expression values so that

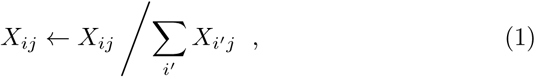

where the left arrow indicates reassignment of the matrix values. Because gene expression values in RNA-seq measurements tend to span many orders of magnitude, it is helpful to apply a standard log_2_ transformation, which is done either to get “fold changes” when comparing groups in differential expression analysis, or to get a “normal” looking statistical distribution. However, the many zeros inherent in single cell RNA-seq data requires the zeros to be replaced with positive values. We choose to replace all zeros with *m*, the smallest nonzero value in *X*_*ij*_, so that

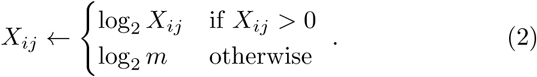

Finally, we keep only those genes exhibiting sufficiently high variation as parameterized by a threshold *θ*,

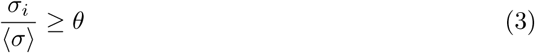

where *σ*_*i*_ is the standard deviation of gene *i*’s expression across all cells and 〈*σ*〉 = *N*^-1^∑_*i*_*σ*_*i*_. For this analysis, we chose *θ* = 0.3.

### Clustering

The clustering algorithms used in p-DCS require to specify the number of clusters *n*. The first step is therefore to find a good value for the parameter *n*. We used the Adjusted Rand Index (ARI) [13] between pairs of clusterings obtained from the same set using a stochastic algorithm (Mini-batch K-Means) and averaging the results to obtain the ARI curve as a function of *n*. The optimal *n* corresponds then to the first peak coming from the *n* = *∞* side of the ARI curve (see Fig. 5 below for an example). To remove noisy components and accelerate the procedure, clustering is conducted on a smaller array 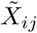 defined by projecting *X*_*ij*_ onto its first 100 principal components (i.e 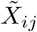 has *i* = 1 *…* 100). The cells in 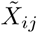 can be clustered using any method available in `scikit-learn` [14] or any custom clustering object with matching syntax. For this application, agglomerative clustering was selected. Clustering diagrams such as Fig. 1(a) are generated by running `scikit-learn`’s t-SNE routine on 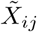 projecting from 100 to two dimensions (simply for the sake of generating a figure). Cells are colored according to their cluster index.

### Cell type assignment

The cell type assignment is based on our voting algorithm idea that uses a database of marker genes. Since this application focuses on bone marrow data, we used Human Cell Markers [1] as our marker/cell type database, *D*. The latter is used to create a marker/cell type table, specific to a gene expression dataset of interest, e.g. the matrix *X* of BM1. The table for a given dataset is created after the initial gene filtering and normalization discussed above. For each cell type in *D* we keep the top *N*_*max*_ = 20 most expressed genes according to an average across all cells in the dataset, thus ensuring that each cell type has at most *N*_*max*_ markers. Additionally, cell types with less that *N*_*min*_ = 4 markers are discarded. The approval votes for each candidate cell type are therefore bounded between *N*_*min*_ and *N*_*max*_. In this way we build a marker/cell type matrix *M*_*km*_ where *k* is the cell type (e.g. T cell), *m* is the marker gene (e.g. CD4). The element *M*_*km*_ = 1 if *m* is a top-*N*_*max*_ most expressed marker of cell type *k* and 0 otherwise.

Building the matrix *M*_*km*_ represents the first step of the voting algorithm. This is equivalent to defining “ballots” in which each qualified voter, i.e. the *N*_*max*_ (or fewer) markers chosen, has a list of candidate cell types they can approve. For each cluster *c*, the voting algorithm is then implemented as follows:

(i) We build the marker/centroid matrix *Y*_*mc*_, where *Y*_*mc*_ is the mean expression of marker *m* across all cells in cluster *c*. For each marker *m*, we use *Y*_*mc*_ to compute all cluster centroids’ z-scores *Z*_*mc*_. Then we build the matrix 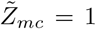 if *Z*_*mc*_*≥ ζ* and 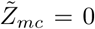 otherwise for a given threshold *ζ*. For this application, we chose *ζ* = 0.3, which provides a reasonable number of markers for all cell types. and This procedure is needed to identify markers that are significantly expressed in one cluster compared to the other clusters. Fig. 3 (a) shows *Y*_*mc*_, calculated for HCA BM1 dataset: darker blue color corresponds to higher expression of markers, and the stars correspond 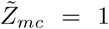, i.e. statistically significant markers with z-score larger than *ζ* among all markers as tested across clusters.
(ii) We compute the vote matrix according to 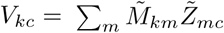. This is when each voter (the markers) matches a given cluster to a single or more possible cell types. This matrix contains an approval score for each type-cluster pair (*k*, *c*).
(iii) To quantify the statistical significance of the approval scores and make the final assignment, we use a stochastic method to quantify the statistical uncertainty associated to each type-cluster pair (*k*, *c*). We create copies of the cells clusters and repeat steps (i) and (ii) *n* = 10^4^ times, each time randomly shuffling cells across clusters. This method accounts for cluster sizes, the overall gene expression distribution of the markers, and imbalances in the number of markers per cell type in estimating the uncertainty. The procedure provides distributions of voting results *𝒫*_*kc*_(*V*_*kc*_) for a null model of random clusters. Fig. 4 (a) shows histograms of the distributions *𝒫*_*kc*_(*V*_*kc*_) calculated for the same dataset of Fig. 3. The figure shows each cell type as a separate plot, and each plot contains the distributions of each cluster in a different color. Note that the distributions do not show a strong dependence on the cluster index *c*, but they can be very different for different cell types *k*.
(iv) Finally, we determine the z-scores, Λ_*kc*_, of the voting results *V*_*kc*_ in (ii), given the null distribution *𝒫*_*kc*_(*V*_*kc*_) calculated in (iii) and assign the cell type according to *T*_*c*_ = argmax_*k*_ Λ_*kc*_. All cells belonging to cluster *c* are thus identified as cell type *T*_*c*_. Fig. 4 (b) is a visual representation of Λ_*kc*_, shown only for positive values, where the indices *k*, *c* are along the x-and y-axis, respectively. After the cell types are determined, the panel (b) of Fig. 3 is produced, with all the markers supporting the assigned identification marked as red stars.

Note that this marker/cell type table is only one of many possible reasonable choices. The software was designed to allow the user to easily substitute this table with a custom table relevant to the particular cell population under investigation. Likewise, the voting scheme outlined above can be replaced with any custom function with the same inputs and outputs. See the documentation for details and examples. [15]

**Figure 3.**
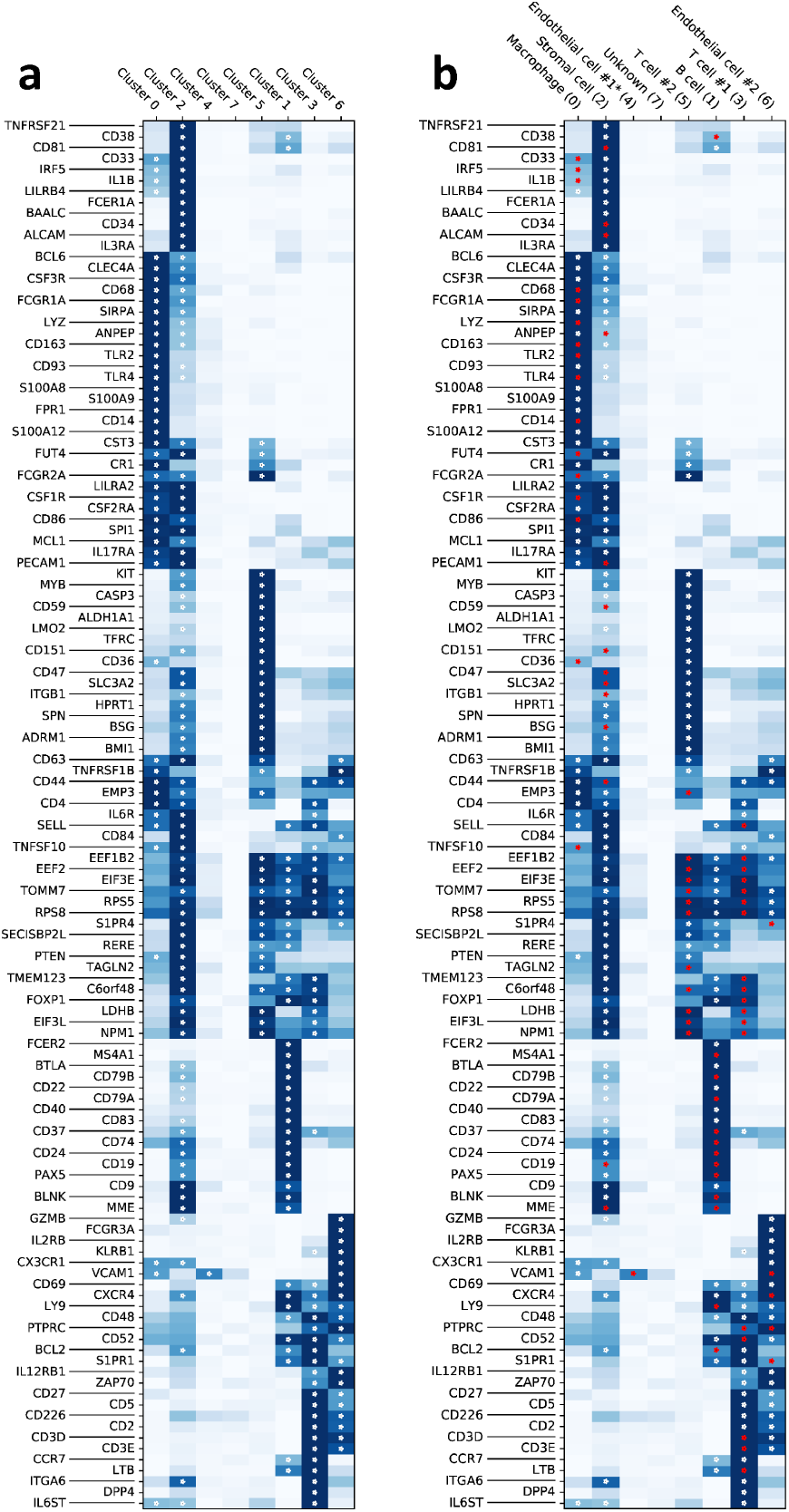
Marker expression for scRNA-seq HCA BM dataset, subset BM1. (a) Mean expression of marker genes in clusters of yet unidentified cell types. Stars denote genes expressed above a certain z-score threshold. (b) Mean expression of marker genes in clusters with inferred cell type with cluster index in parentheses. Red stars highlight the supporting markers in assigning the cluster cell type.

**Figure 4.**
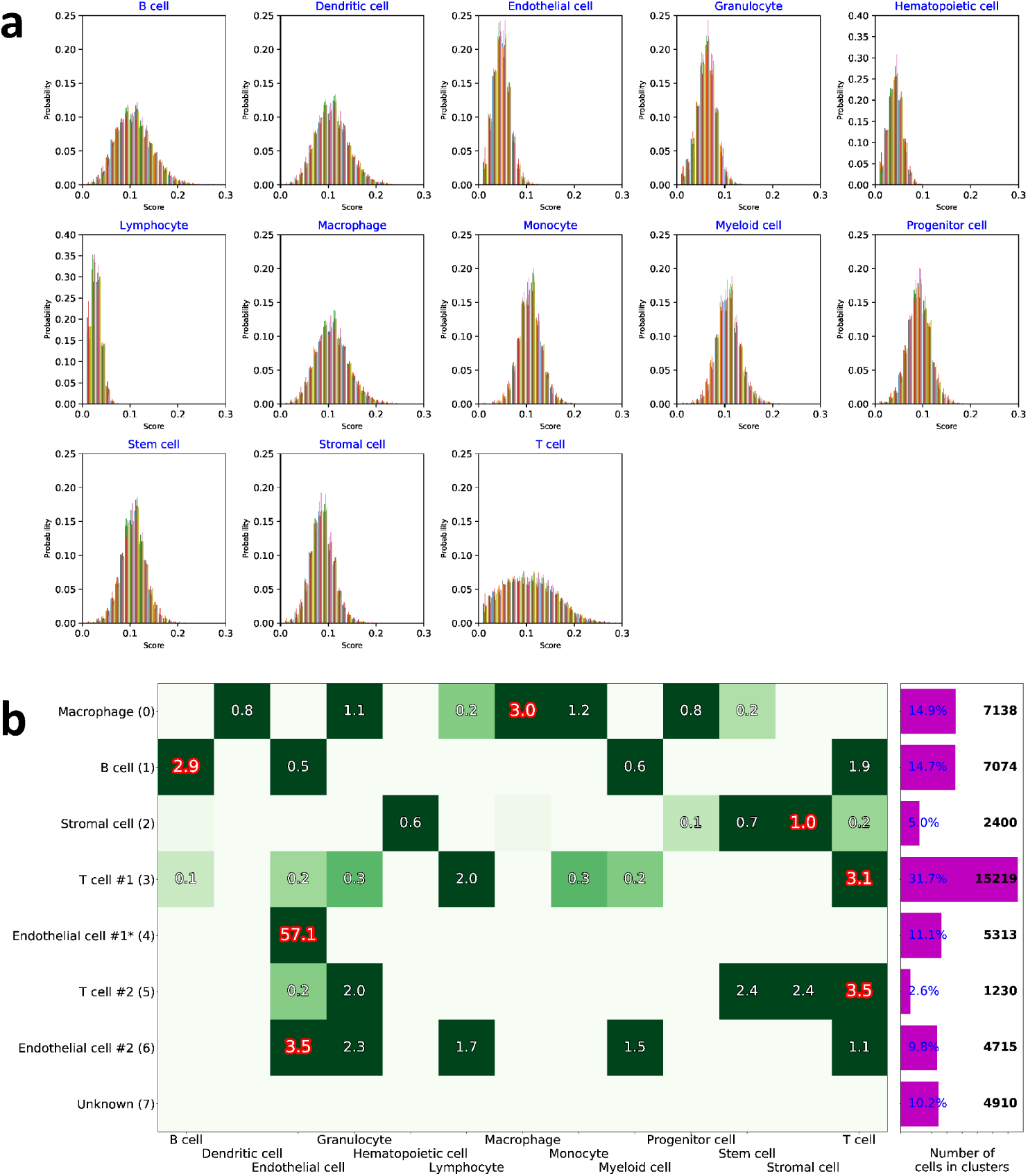
Voting results visualization. Exemplified on HCA BM1 dataset. (a) *𝒫*_*kc*_(*V*_*kc*_) distributions shown in separate plots for each cell type *k*, different cluster *c* are shown in different color. (b) Visualization of the matrix Λ_*kc*_, where columns are the possible cell types and rows are the assigned cell types *T*_*c*_, with cluster indices 0,1,…,7 in parentheses. The negative z-scores are not shown. The barplot on the right shows relative (%) and absolute (cell count) cluster sizes. Cell clusters that have 3 or less supporting markers are marked with “*”, see Figure 2 for supporting markers.

## Results and discussion

In this section, we first present the results obtained with our methodology using recently-published data from normal bone marrow samples (the data identified above as BM1-BM8, containing a total of 378k cells). Additionally, we compare our cell type assignment to an existing identification of cell types from a large scRNA-seq ∼68.6k cells PBMC dataset.

### Results on the HCA BM data

#### Number of clusters

We first calculated the Adjusted Rand Index (ARI) [13] curves for BM1-BM8. For each *n* between 4 and 16, Mini-batch K-Means clustering was performed 12 times leading to 12 different partitions of the data. The ARI between all the possible 66 pairs of partitions was then calculated and averaged. The procedure was repeated in *N* = 200 independent runs to obtain error bars. The ARI curves are shown in Fig. 5. Note that the ARI curves often have a maximum at or near *n* = 1. This maximum does not provide useful information, and the optimal *n* is therefore associated to the first peak observed coming from the right side of the plot. In addition to the ARI for each of the BM1-BM8 sets, Fig. 5 displays their average in black. The latter has a peak at *n* = 8, and we therefore select that value for clustering all the datasets.

#### Clustering and identification in BM1-BM8 datasets

The BM samples were analyzed individually and their cluster plots were combined to demonstrate the similarity between the 8 datasets of bone marrow, see Fig. 6. We restricted the candidate cell types to the ones that have more than three markers expressed in each dataset after pre-processing. The color coding is uniform for the cell types across the 8 datasets, i.e. all Stromal cells are colored orange, B cells – dark blue, etc. As some of the clusters overlap on the t-SNE plot [16, 17], it is useful to calculate the relative fractions of cells of various cell types. The latter provide a snapshot of the cellular composition of the 8 bone marrow samples, see Fig. 7.

**Figure 5.**
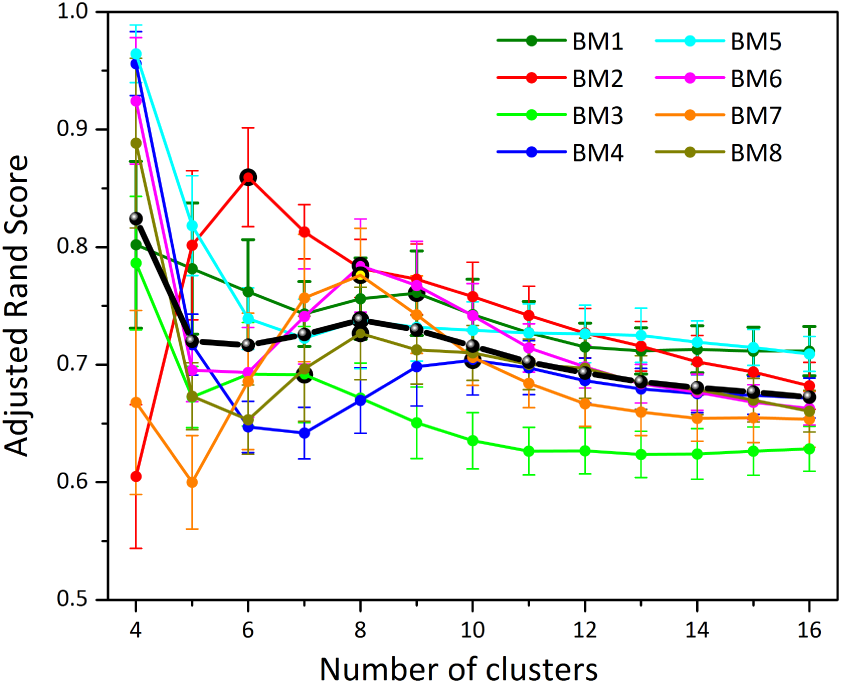
HCA BM dataset analysis. Adjusted Rand Index (ARI) curves for each dataset BM1-BM8. Clustering was done using Mini-Batch K-Means from scikit-learn. The black line represents the average of the 8 datasets, and the peak at *n* = 8 was used to select the optimal number of clusters.

**Figure 6.**
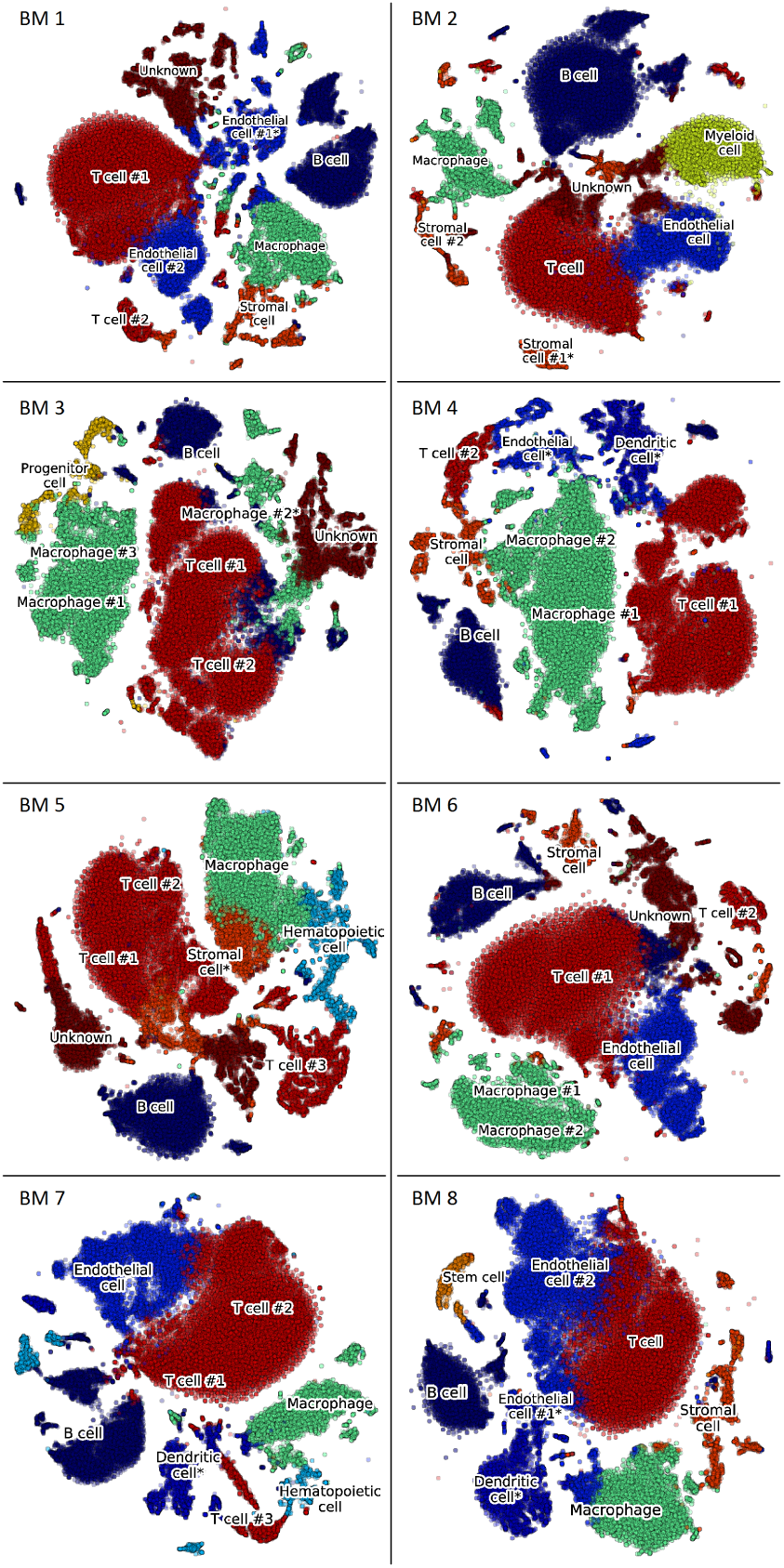
HCA BM preview dataset analysis. Clustering illustrated with t-SNE plots for each patient in the dataset. The cell type identification is assigned based on the voting algorithm discussed in Methods.

**Figure 7.**
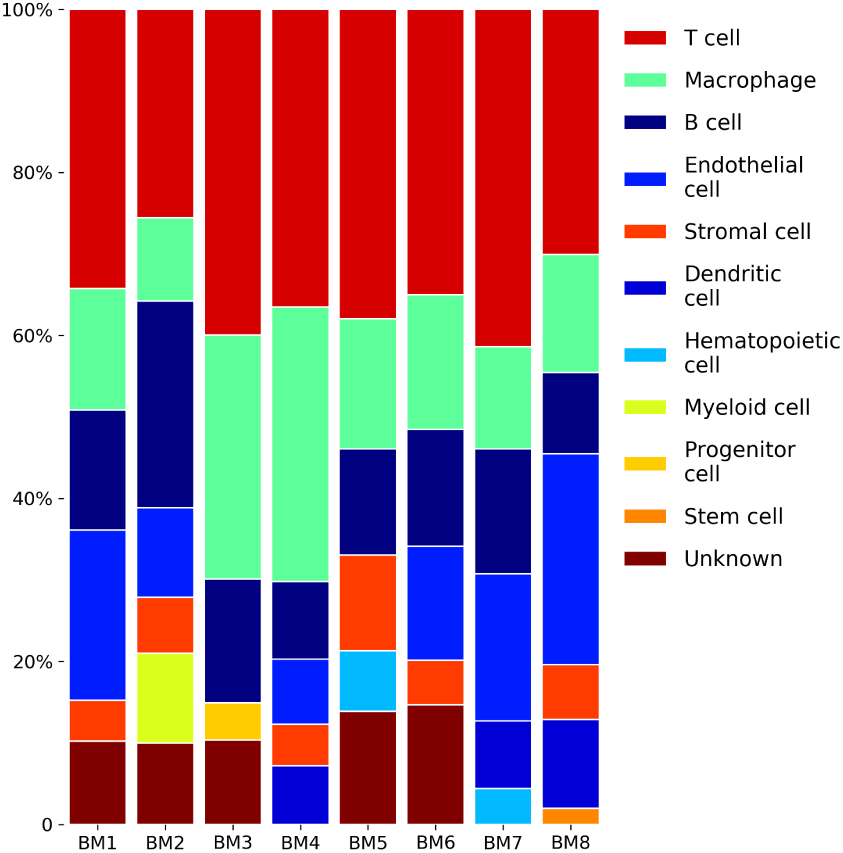
HCA BM dataset summary. Cell type relative fractions for each BM sample. The cell types are sorted by average (across samples) fraction size, with the exception of the “Unknown” which is moved to the bottom. Color coding for cell types is identical to Fig. 6.

#### Clustering of T and B cells sub-types

We applied the methodology illustrated above to identify sub-types of major hematological B and T cells. Additional marker/cell subtype tables *M*_*km*_ were prepared for this analysis. Columns of these new matrices indicates sub-types only and rows are the markers/genes that are known to be expressed the these sub-types. We used the same Human Cell Markers [1] database to build the *M*_*km*_ matrices for B and T cells. As above, these matrices *M*_*km*_ are created ensuring that only the top *N*_*max*_ = 20 most expressed makers are included for each sub-type. Cell sub-types with no expressed makers after pre-proceessing are discarded.

Clustering with *n* = 5 for T cell subtypes from BM1 is shown in Fig. 8 (a), revealing Nave T cell and Helper T subtypes. In the same way, B cells of BM1 were processed into 5 clusters in Fig. 8 (b), showing populations of Transitional T1 and T2 B cells and a small group of Plasma cells.

**Figure 8.**
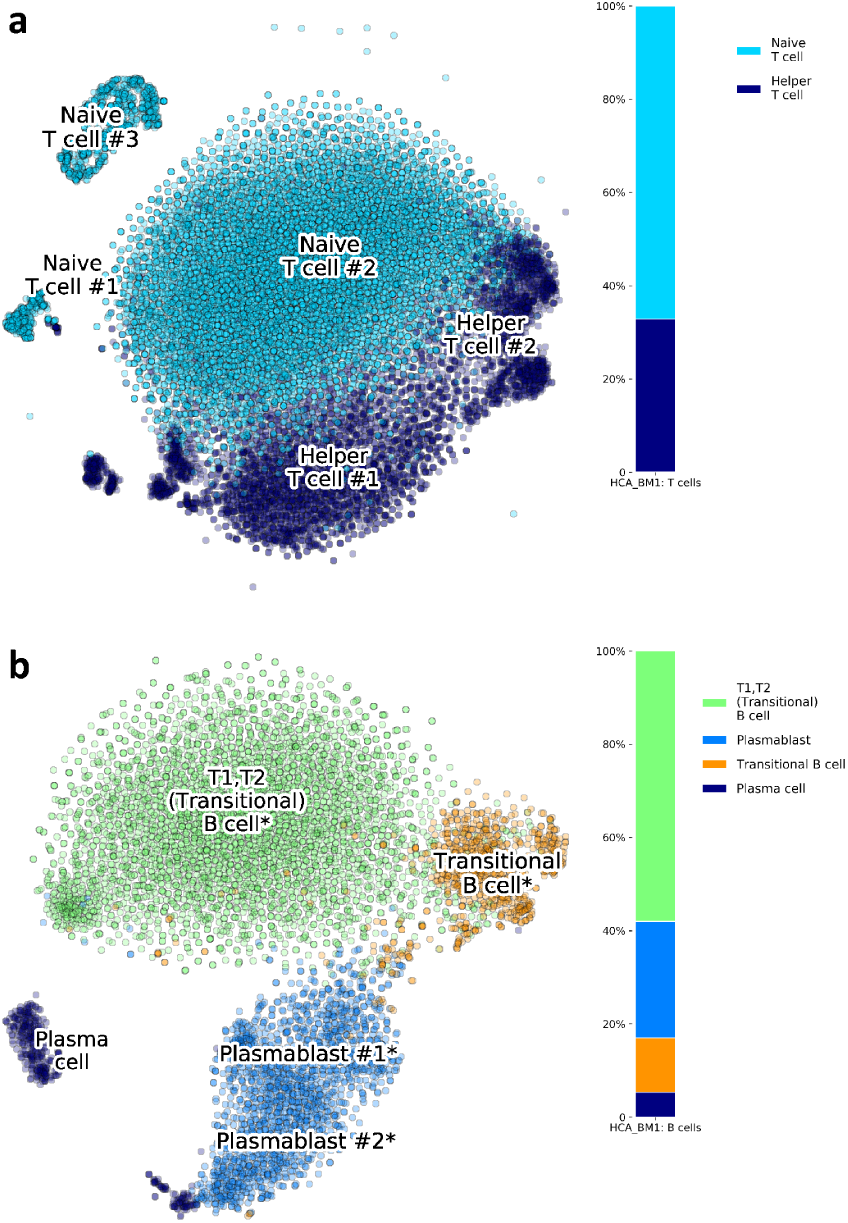
Subclustering of HCA BM1. Application of p-DCS on (a) T cells, and (b) B cells, revealing subtype composition.

Note that the simultaneous identification of major cell types and their relative sub-types is problematic. The best approach consists in first identifying major cell types and then separately analyzing each of them as shown in this section. We have tried to include major cell types and their subtypes in the matrices *M*_*km*_ and have attempted their identification with a larger number of clusters. Such an approach leads often to incorrect results with relative cell frequencies that are incompatible with normal physiological ranges.

#### Congruence with expert annotation on PBMC dataset

In a recent work, Sinha et al. [9] presented their dropClust algorithm to cluster ultra-large scRNA-seq datasets. To illustrate their algorithm, they used data from 68k PBMC from Zheng et al. [4]. Their cluster annotation, obtained from a manual assessment using a few selected markers, is of interest here and can be used to compare the annotation obtained by our automated methodology with one obtained manually by an expert. By pre-processing the whole 68k PBMC dataset, we determined that the optimal number of clusters was 8. The result of the analysis is shown in Fig. 9. The clustering and cell type inference from the automated p-DCS procedure are shown in Fig. 9 (a), indicating that T cells constitute the major cell type in this sample. Fig. 9 (b) shows a graphical comparison of cell types fractions obtained by p-DCS and by Sinha et al. [9]. The frequencies of various cell types are expected to vary from individual to individual, and the fractions that we determined are within the normal ranges [18]. The main difference in cell type frequencies, Fig. 9 (b), determined using two approaches is in p-DCS NK cell cluster (yellow) which in Sinha et al. is split into NK (yellow) and NK T (light blue) cells. The latter cell type expresses a combination of T cell and NK cell attributes and markers and therefore categorizing NK and NK T cells is challenging. Fig. 9 (c) displays the cell types used in voting and z-scores of the voting scores. The quantitative comparison is also available in Table 1. In addition to comparison of sizes of cluster between the two methods, p-DCS and dropClust, we individually analyzed all cells, i.e. their barcodes in the scRNA-seq data, to check if they were assigned to matching cell types. For each cell type annotated by p-DCS we counted how many cells were annotated by Sinha et al. [9] into each of their categories (Table 2). Overall the agreement is strong, with the exception of Dendritic cells and Macrophages for which we observed a significant mismatch.

**Table 1.** Comparison of p-DCS and DropClust on PBMC scRNA-seq ∼68.6k cells dataset.

| *Cell type* | **p-DCS** | **DropClust** |
| --- | --- | --- |
| T cell | Cluster #1, 4, 6: 72.2% | Cluster #1, 2, 5, 7, 10: 73.2% |
| NK T cell |  | Cluster #3: 9.0% |
| NK cell | Cluster #0: 14.4% | Cluster #6, 13: 5.0% |
| B cell | Cluster #3: 5.7% | Cluster #4: 5.8% |
| Dendritic cell | Cluster #5: 3.0% | Cluster #11, 14: 1.8% |
| Macrophage | Cluster #2: 4.4% |  |
| Monocyte |  | Cluster #8, 9: 4.9% |
| Megakaryocyte | Cluster #7: 0.2% |  |
| Circulating Megakaryocyte Progenitors |  | Cluster #12: 0.2% |

**Table 2.** Cell counts from cell-by-cell validation of p-DCS and dropClust on PBMC scRNA-seq ∼68.6k cells dataset.

| *p-DCS cell type (count)* | *dropClust cell type* |  |  |  |  |  |  |
| --- | --- | --- | --- | --- | --- | --- | --- |
|  | *T cell* | *NK T cell* | *NK cell* | *B cell* | *Monocyte* | *Dendritic cell* | *CMP\** |
| T cell (49559) | 48951 | 494 | 69 | 42 |  | 1 | 2 |
| NK cell (9894) | 914 | 5681 | 3298 | 1 |  |  |  |
| B cell (3931) | 84 | 5 | 1 | 3841 |  |  |  |
| Macrophage (2990) | 57 | 7 |  | 76 | 1802 | 1048 |  |
| Dendritic cell (2042) | 235 | 13 | 1 | 19 | 1537 | 237 |  |
| Megakaryocyte (163) |  |  |  |  |  |  | 163 |

\*CMP–Circulating Megakaryocyte Progenitor

**Figure 9.**
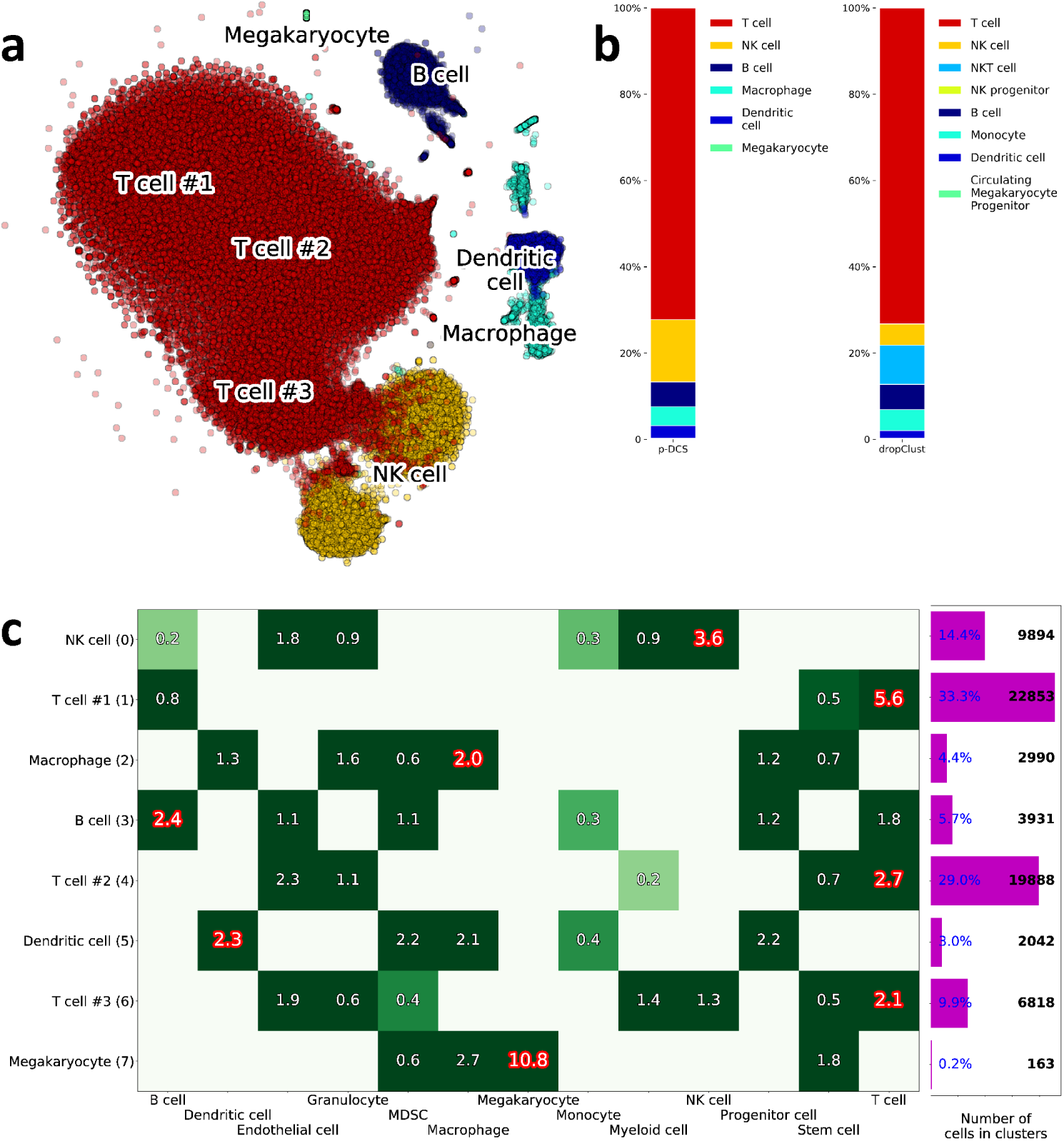
PBMC dataset processing. (a) Clustering with inferred cell types. (b) Fractions of various cell types obtained by p-DCS in comparison with DropClust manual clusters’ annotations [9]. (c) Visualization of the voting results of all possible cell types (columns) and identified clusters (rows), generated from the input marker cell/type table.

Sub-clustering of T cells was also done to compare the two approaches. T-cells from clusters 1, 4 and 6 (see Fig. 9) were processed with a new list of markers/cell sub-types. The results of cell sub-types annotation are presented in Fig. 10, and the detailed comparison to the results by Sinha et al. [9] are in Table 3.

**Table 3.** Sub-clustering of ∼49.6k T cells subset of ∼68.6k cells dataset. Comparison of p-DCS and DropClust subtypes assignment.

| *T cell subtype* | **p-DCS** | **DropClust** |
| --- | --- | --- |
| Naive T cell | Cluster #0, 6, 7: 41.7% | Cluster #1: 46.0% |
| Cytotoxic T cell (CD8+ T cell) | Cluster #2.5: 13.7% | Cluster #5,7: 11.8% |
| T helper cell | Cluster #1: 12.0% |  |
| Unknown | Cluster #3: 4.3% |  |
| Memory T cell |  | Cluster #2: 14.9% |
| Regulatory T (Treg) cell | Cluster #4: 1.6% | Cluster #10: 0.5% |

**Figure 10.**
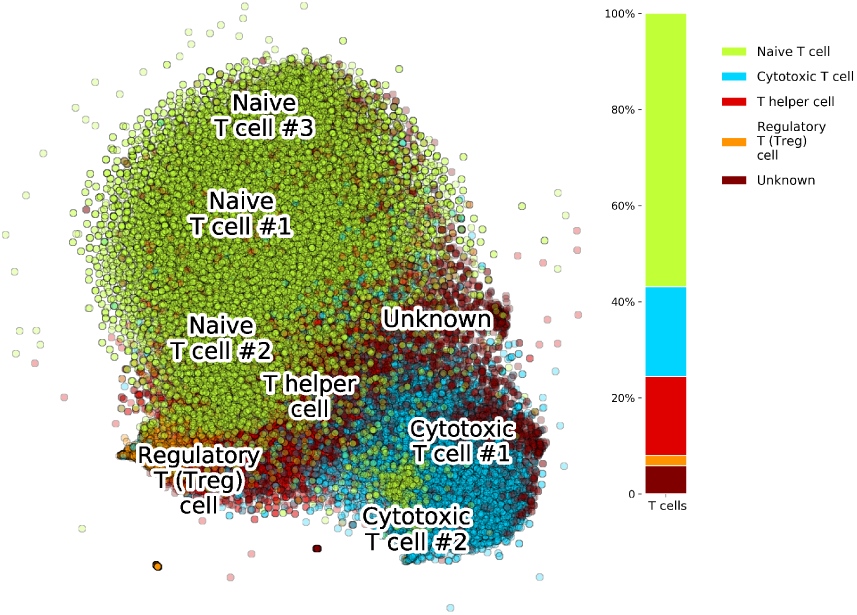
PBMC dataset T cells subset analysis. Sub-clustering of cells from clusters 1, 4 and 6 of the PBMC dataset reveals that the p-DCS automatic sub-type identification is in good agreement with manual annotation.

### Alternative cell marker input lists

We have used the cell marker database CellMarker [1], which is the most recent database available. There is another database created by the Human Cell Differentiation Molecules (HCDM) organization [19], which is sponsored by a number of large companies. This database contains detailed information about each CD molecule, including structure, function, and cellular expression. The HCDM would be an alternative to CellMarker, that could be used to create a marker/cell type table to employ with p-DCS. We have observed that the overlap between these two databases is very strong, therefore we do not expect significant differences in the cell cluster assignments. Finally, several deconvolution algorithms have been developed in the past for estimating the relative composition of complex tissues from bulk transcriptomics data. [20, 21, 22, 23, 24, 25, 26, 27] These methodologies are usually based on predefined signature matrices that contain the relative expression of markers, not just the presence/absence of a marker, for different cell types. Regression methods are then typically used to infer the relative proportions in a mixture. These signature matrices have been validated on bulk data and their robustness to the characteristic scRNA-seq noise has not been tested. However, in principle they contain additional information that could be integrated in our p-DCS to identify single cells.

## Conclusions

We have presented a methodology that, after unsupervised clustering of scRNA-seq data, automatically assigns clusters to cell types based on a voting algorithm without manual interpretation by an expert curator. The method provides the classification of individual cells into predefined classes based on a comprehensive database of known molecular signatures, i.e. cell surface (extracellular) and intracellular markers [1]. The proposed methodology assures that extensive marker/cell type information is taken into account in a systematic way when assigning clusters to cell types. Moreover, the method allows for a high throughput processing of multiple scRNA-seq datasets since it does not involve an expert curator.

In addition to determining major cell types, we have shown how this methodology can be applied recursively to obtain cell sub-types. We have performed a congruence analysis of cluster identification obtained by our method with those obtained by expert curators on the same dataset, showing that the automatic assignment is consistent with expert assignment both of major cell types and cell sub-types. While we have focused on the identification of hematological cell types, the software is designed to allow the user to substitute the marker table to apply the methodology to different tissues.

## Abbreviations

ARI: Adjusted Rand Index
BMMC: Bone marrow mono-nuclear cells
CD: Clusters of Differentiation
HCA: Human Cell Atlas
HCDM: Human Cell Differentiation Molecules
PBMC: Peripheral blood mono-nuclear cells
PCA: Principal Component Analysis
p-DCS: Polled Digital Cell Sorter
tSNE: t-distributed Stochastic Neighbor Embedding

## Acknowledgements

We thank Prof. George I. Mias and Prof. Michael Bachmann for helpful suggestions.

## Funding

This work was supported by National Institutes of Health, Grant No. R01GM122085.

## Availability of data and materials

Analyzed here HCA BM data, available to the research community, was obtained from HCA Data Portal https://preview.data.humancellatlas.org/. The 68k PBMC data, by Zheng et al., used for the p-DCS methodology validation is available at https://support.10xgenomics.com/single-cell-gene-expression/datasets/. The software is available as a python package at https://github.com/sdomanskyi/DigitalCellSorter.

## Author’s contributions

SD, AS, and CP designed the algorithms. SD, AS, JW, and NH wrote the software. GP provided bio-medical analysis. SD, AS, and CP wrote the manuscript.

## Ethics approval and consent to participate

Not applicable

## Consent for publication

Not applicable

## Competing interests

JW is an employee of Salgomed Inc., and CP and GP own equity in Salgomed Inc.

